# 6S RNA supports recovery from nitrogen depletion in *Synechocystis* sp. PCC 6803

**DOI:** 10.1101/134676

**Authors:** Beate Heilmann, Kaisa Hakkila, Jens Georg, Taina Tyystjärvi, Wolfgang R. Hess, Ilka M. Axmann, Dennis Dienst

## Abstract

**Background:** The 6S RNA is a global transcriptional riboregulator, which is exceptionally widespread among most bacterial phyla. While its role is well-characterized in some heterotrophic bacteria, we subjected a cyanobacterial homolog to functional analysis, thereby extending the scope of 6S RNA action to the special challenges of photoautotrophic lifestyles.

**Results:** Physiological characterization of a 6S RNA deletion strain (Δ*ssaA*) demonstrates a delay in the recovery from nitrogen starvation. Significantly decelerated phycobilisome reassembly and glycogen degradation are accompanied with reduced photosynthetic activity compared to the wild type.

Transcriptome profiling further revealed that predominantly genes encoding photosystem components, ATP synthase, phycobilisomes and ribosomal proteins were negatively affected in Δ*ssaA*. *In vivo* pull-down studies of the RNA polymerase complex indicated a promoting effect of 6S RNA on the recruitment of the cyanobacterial housekeeping σ factor SigA, concurrently supporting dissociation of group 2 σ factors during recovery from nitrogen starvation.

**Conclusions:** This study reveals 6S RNA as an integral part of the cellular response of *Synechocystis* sp. PCC 6803 to changing nitrogen availability. According to these results, 6S RNA supports a rapid acclimation to changing nitrogen supply by regulating the switch from group 2 σ factors SigB, SigC and SigE to SigA-dependent transcription.

## Background

Cyanobacteria are photoautotrophic prokaryotes essentially relying on the availability of sunlight and CO_2_ as their major energy and carbon source, respectively. Due to their autotrophic lifestyle and versatile metabolism, cyanobacteria are suited as economical cellular chassis for diverse biotechnological applications like energy feedstock accumulation [1], third generation biofuel production [2-5] and commodity product biosynthesis [6-8]. However, the long-term mass cultivation for these applications at high cell densities also causes stress conditions that arise from redirecting metabolism, accumulation of potentially toxic compounds and the concomitant tendency for nutrient limitation (reviewed by Dexter *et al*. [2]). Hence, the requirements for molecular tools and well-defined targets for manipulating the cellular program towards optimized nutrient utilization are of growing interest. Within the diversity of their natural terrestrial and aquatic habitats, cyanobacteria have certainly developed extensive regulatory systems to acclimate to nutrient and light limiting conditions, which are also involving small regulatory RNA molecules (sRNAs, e.g. [9-14]).

One of the best-characterized prokaryotic sRNA regulators is the widely conserved 6S RNA that responds to changes of the nutritional status [15-18]. When cells of *Escherichia coli* (*E. coli*) enter the stationary growth phase, this highly structured RNA specifically regulates transcription by promoter mimicry, as the RNA polymerase (RNAP) holoenzyme carrying the housekeeping sigma factor σ^70^ binds to 6S RNA instead of the promoter regions of the household genes [19-22]. Upon nutrient-induced outgrowth from the stationary phase, which includes enhancement of NTP levels, 6S RNA acts as a template for the de novo synthesis of a ∼20 nt product RNA (pRNA) [23, 24]. pRNA synthesis triggers the release of 6S RNA from RNAP reverting 6S RNA-dependent inhibition [23, 25]. 6S RNA lacking mutants of *E. coli* and *Bacillus subtilis* (*B. subtilis*) show significant phenotypes under long-term nutrient deprivation [20], stress conditions, like alkaline stress [29, 30] and during outgrowth from stationary phase [31, 32]. Furthermore, the presence of 6S RNA leads to an upregulation of σ^38^-activity [33, 34], while overexpression of 6S RNA in σ^38^-deficient cells results in reduced viability in late stationary phase [19].

Homologues of 6S RNA have been identified in several freshwater cyanobacteria, including the model organism *Synechocystis* sp. PCC 6803 (*Synechocystis* 6803) [15, 26, 27]. Whereas *in vitro* studies indicate a conservation of basic 6S RNA mechanisms, little is known about the functional relevance of the cyanobacterial 6S RNA *in* vivo [28].

Since phototrophic growth of cyanobacteria does essentially rely on atmospheric or enriched CO_2_ supplementation and – importantly – light, plain dilution experiments would rather evoke responses to changing light availability (and its relation to CO_2_ availability). Accordingly, a specified experimental setup was required for functional characterization of cyanobacterial 6S RNA during outgrowth from nutritional deprivation. Regarding the status of an inorganic nitrogen source – i.e. nitrate or ammonia – non-diazotrophic cyanobacteria like *Synechocystis* 6803 exhibit a well-controlled acclimation behavior that is also accompanied by a stationary growth plateau [35].

Acclimation of cyanobacteria to changing nitrogen availability is a tightly controlled process that has been well described in the literature. One of the most pronounced physiological responses is the remodeling of the photosynthetic machinery, leading to distinctly and visibly reduced amounts of the phycobilisome antenna complex and – delayed in time – chlorophyll *a* breakdown [35-38]. These biochemical mechanisms are influenced by elaborate underlying events of gene expression control. This network essentially involves the collaborative action of the global transcriptional regulator NtcA and the proteins PII and PipX [39].

Moreover, several group 2 σ factors appear to play a role within the nitrogen regulatory network. Besides the nitrogen stress response σ factor SigE, these are SigB, involved in the NtcA-dependent nitrogen-related gene expression during exponential growth and SigC, in the response of stationary phase cultures to nitrogen starvation [40-42].

Several reports described the transcriptomic characteristics of nitrogen-starving cyanobacteria [43, 44] as well as the response upon nitrogen repletion in a time-resolved manner [38, 45]. The recovery process after long-term starvation is initiated by the upshift of respiratory gene expression and switching on the translational machinery as well as nitrogen assimilation.

Further, until full physiological restoration, the cells successively reactivate photosynthetic complexes, carboxysomes and the cell division machinery [45]. Cells without any group 2 σ factors are not able to recover from nitrogen deficiency [42].

Here we describe the involvement of the sRNA regulator 6S RNA in the dynamics of nitrogen acclimation in *Synechocystis* 6803, elaborating the example of NO_3_^-^-mediated recovery from long-term nitrogen starvation.

## Results

### Inactivation of 6S RNA affects photosynthetic parameters and storage of carbohydrates upon nitrogen starvation and recovery

To analyze the functional role of 6S RNA in cyanobacteria, we deleted the *ssaA* gene encoding 6S RNA from the genome of *Synechocystis* 6803. The *ssaA* gene was replaced with a kanamycin resistance cassette (Fig. 1a) and complete segregation of the resulting Δ*ssaA* mutant was confirmed by colony PCR (Figure 1b) as well as by Northern Blot analysis (Fig. 1c).

**Figure 1.**
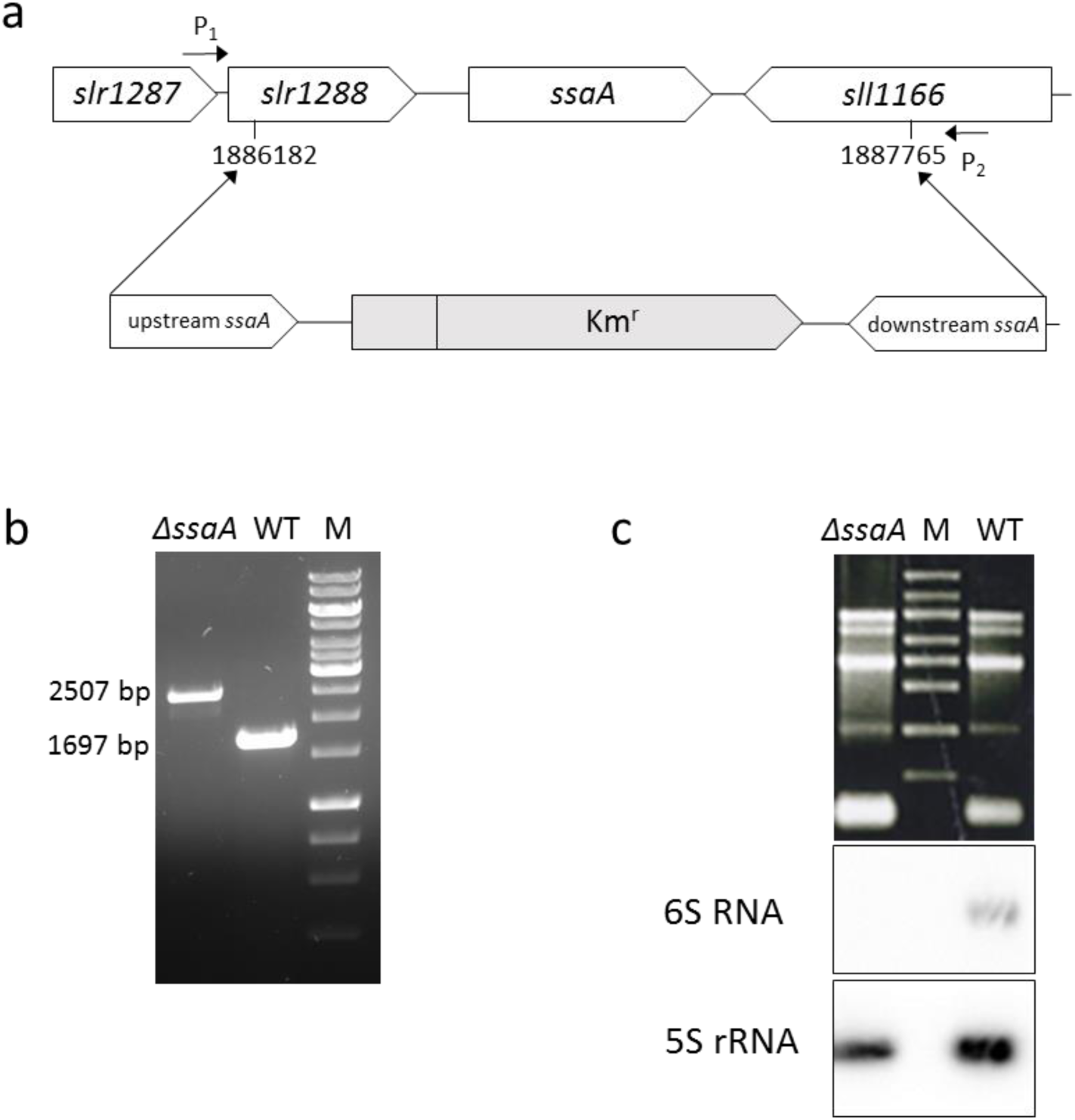
Strategy and confirmation of the *Synechocystis* 6803 Δ*ssaA* mutant. (a) Strategy for the construction of the Δ*ssaA* mutant. The *ssaA* gene was replaced by a kanamycin resistance cassette (Km^r^) and its promoter region (grey boxes) and flanking regions of ∼ 700 bp upstream and downstream from the *ssaA* gene at the indicated genome positions. P1 and P2 represent the location of primers used to verify complete segregation of the mutant strain. The location of genes *slr1287, slr1288* and *sll1166* are depicted upstream and downstream of the *ssaA* gene. (b) Complete segregation of the Δ*ssaA* mutant gene copies and replacement with a kanamycin resistance cassette (Km^r^) and its flanking regions (998 nt) was tested by PCR analysis, using primers P1 and P2. (c) Deletion of 6S RNA in Δ*ssaA* mutant was confirmed by Northern blot analysis using a radioactive [^32^P]-labelled oligonucleotide for 6S RNA. 5S rRNA was hybridized as a loading control.

Under standard growth conditions, the Δ*ssaA* strain did not show any clear phenotype. Referring to 6S RNA function in *E. coli* and *B. subtilis*, where it regulates RNAP activity during the stationary growth phase and outgrowth [17, 18, 46], the following mutant characterization experiments were conducted under nitrogen depletion (which leads to a stationary state of cell culture) and subsequent recovery by repletion with 17.6 mM NaNO_3_ (corresponding to outgrowth).

The acclimation to nitrogen depletion proceeded similarly in the Δ*ssaA* strain as in wild type (WT), demonstrated by similar whole-cell absorption spectra of Δ*ssaA* and WT cells after seven days of nitrogen deficiency (Fig 2a, 0h +N samples). Furthermore, light-saturated photosynthetic activities (Fig. S1a) and 77K fluorescence spectra (Fig. S1b) were similar in both strains in the middle of the nitrogen deficiency treatment.

Visible differences between the strains were detected during the recovery phase. After 22h of recovery, WT showed normal green color of a viable *Synechocystis* 6803 culture while the Δ*ssaA* mutant still remained yellow-orange, which is a typical characteristic of nitrogen depleted cultures (Fig. 2a). The Δ*ssaA* complementation strain (Δ*ssaA*-c) recovered from the nitrogen deficiency similarly as WT (data not shown).

**Figure 2.**
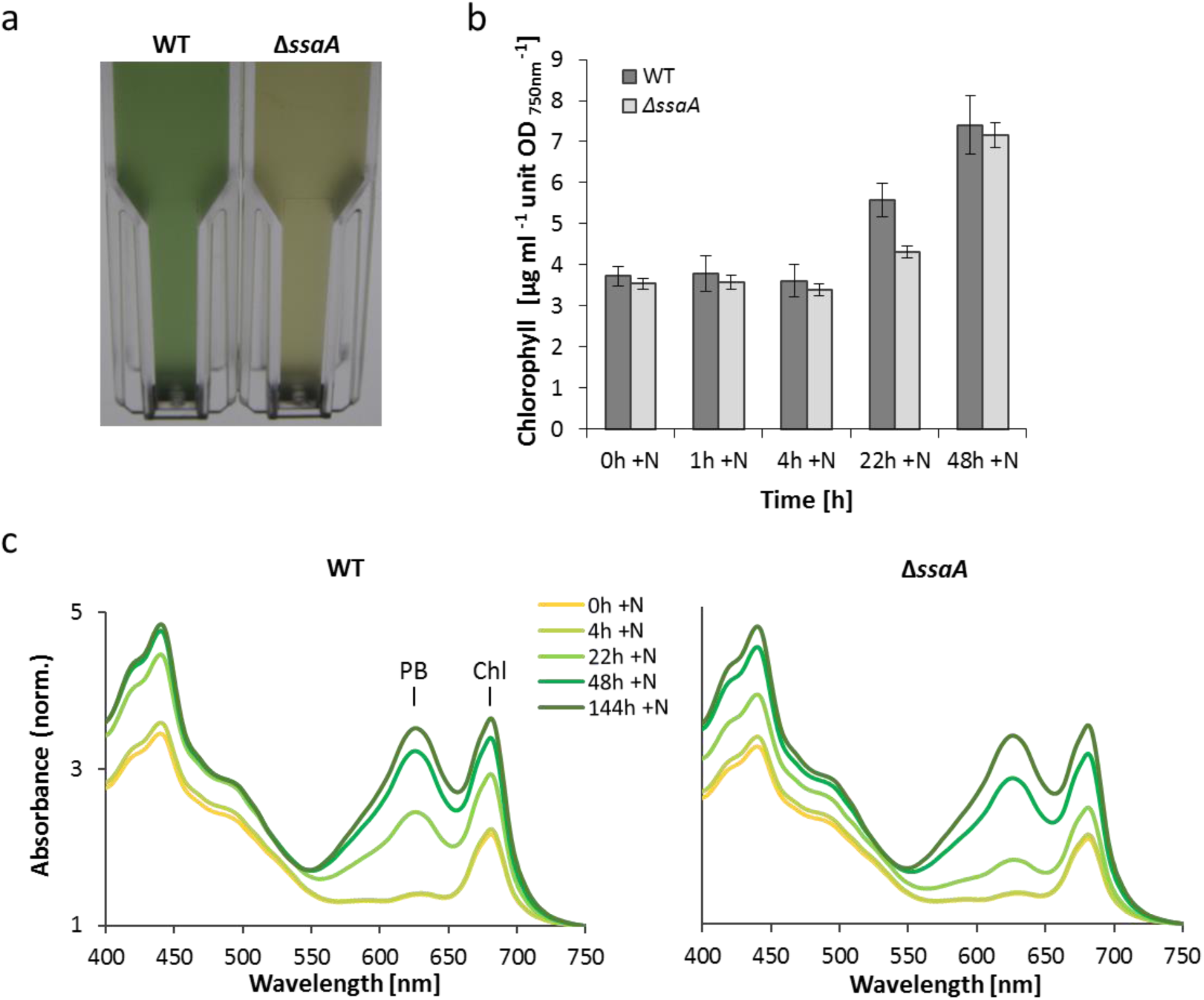
Comparative physiological characterization of the *Synechocystis* 6803 WT and Δ*ssaA* mutant strain and phenotype studies of the mutant strain. Cultures were grown under continuous white light at an irradiance of 80 μmol photons m^-2^ s^-1^, at 30 °C, in nitrogen-depleted medium for 189 hours before recovery was initiated by adding nitrogen. (a) Photograph of cultures of *Synechocystis* 6803WT and Δ*ssaA* mutant strain during recovery from nitrogen starvation, t= 22h +N. Cultures were adjusted so that their cell densities were the same (as estimated by the OD_750_), transferred to 1-cm cuvettes, and photographed with illumination from behind. (b) Chlorophyll content of *Synechocystis* 6803 WT and Δ*ssaA* measured after cultivation in nitrogen deficiency for 189h (t= 0h +N) and at recovery time points t= 1h, t= 4h, t= 22h and t= 48h (+N). The illustrated data represent the mean from three independent biological replicates and the standard deviation was calculated. (c) Details of whole cell absorbance spectra of *Synechocystis* 6803 WT and Δ*ssaA* strains are shown for the time point t= 0h +N, which corresponds to 189h under nitrogen deficiency (-N) and for the time points t= 4, t= 22, t= 48 and t= 144 hours after nitrogen addition (+N). The spectra were normalized at 750 nm. PB: phycobilin; Chl: chlorophyll *a.*

The color of nitrogen depleted cells is due to prominent loss of blue phycobilins (absorption maxima at 620 - 635 nm), clear loss of green chlorophyll *a* (Chl *a*, absorption maxima at 680 nm), but only minor reduction of yellow-orange carotenoid pigments (Fig. 2c; compare 0h +N and the fully recovered 144h +N samples). Re-synthesis of pigments takes hours after nitrogen supply. In both strains phycobilin (Fig. 2c) and Chl *a* (Fig. 2b and 2c) contents remained low after 4h of recovery.

After 22h both phycobilins and Chl *a* showed significant retrieval in WT while in Δ*ssaA,* the recovery process was still in an early stage. A significant delay in phycobilisome reconstruction in Δ*ssaA* was also seen as reduced contents of the main phycobilisome proteins, phycocyanin and allophycocyanin, after 22h of nitrogen repletion (Fig. S2a and c). However, Δ*ssaA* cells completely recovered over extended periods of 48h or longer (Fig. 2b and 2c).

77K fluorescence spectra, measured with orange light phycobilisome excitation 14h after nitrogen addition, showed a lower phycobilisome peak in Δ*ssaA* cells than in WT confirming the slow recovery of phycobilisome antenna in the mutant (Fig. 3a). In addition, the ratio of the photosystem II (PSII) CP47 peak to CP43 peak was lower in the Δ*ssaA* mutant than in WT, indicating reduced efficiency of electron transfer within PSII in Δ*ssaA* (Fig. 3a). In accordance with that the light-saturated PSII activity of the Δ*ssaA* strain was 20% lower than that of WT after 22h of recovery in nitrogen replete conditions (Fig. 3b), and also the light saturated photosynthesis rate of the mutant strain was lower than that of WT (Fig. 3c). According to these results, both light-harvesting capacity and photosynthetic activity recovered more slowly in the mutant than in WT.

**Figure 3.**
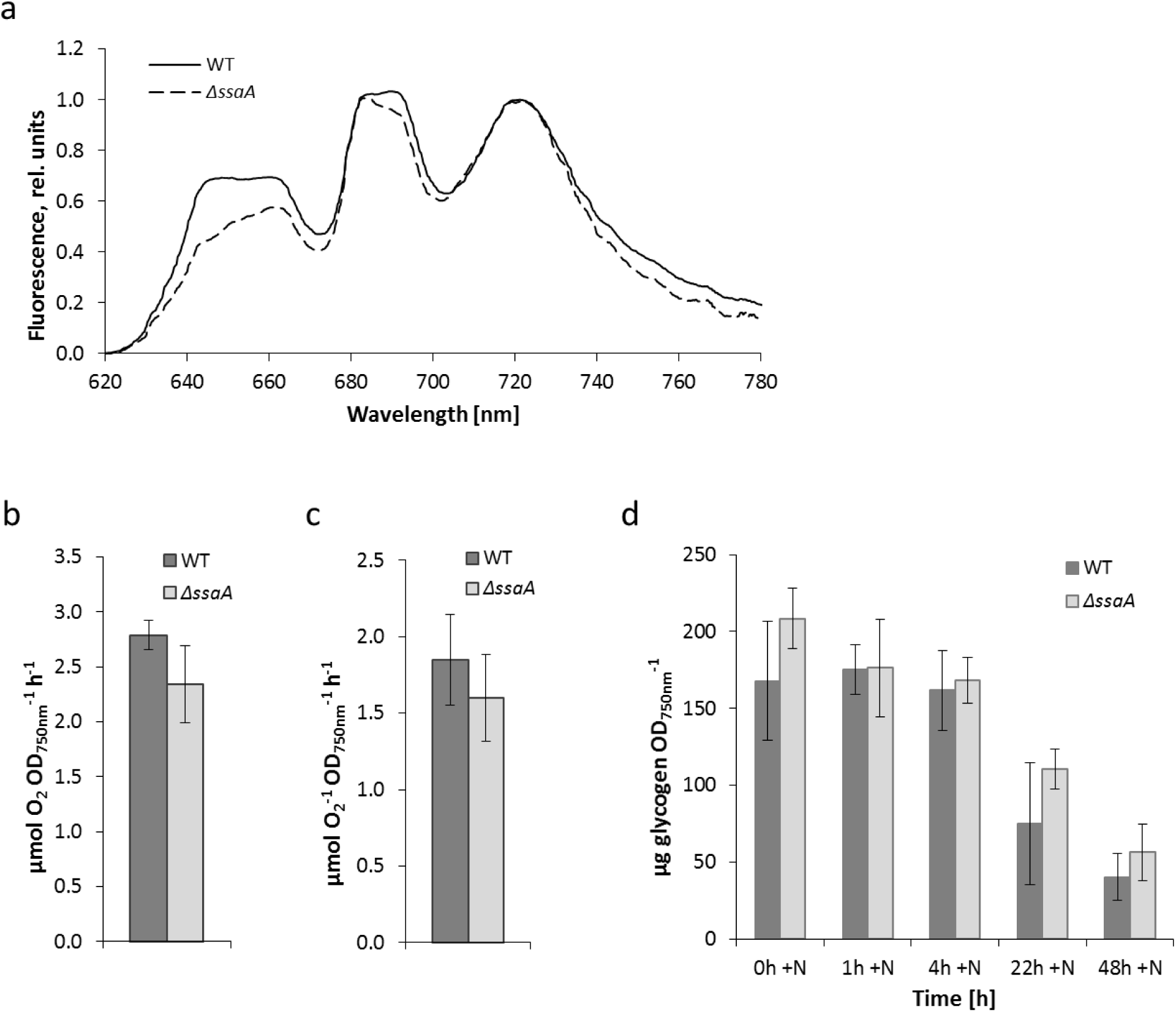
Photosynthetic parameters of WT and Δ*ssaA* mutant and glycogen consumption during recovery from nitrogen starvation. (a) Orange light-excited fluorescence emission spectra of WT and Δ*ssaA* mutant at 77 K after 14 hours of recovery (+N). The spectra were normalized by dividing by the peak value of PSI at 721 nm and setting this value to 1. Oxygen evolution rates of electron transport in PSII (b) and photosynthesis (c) measured *in vivo* at time point 16 - 22 hours +N under light-saturated conditions (3000 µmol photons m^-2^ s^-1^). (d) Glycogen content of *Synechocystis* 6803 WT and Δ*ssaA* after cultivation in nitrogen-depleted medium for 189h (t= 0h +N) and in NaNO_3_-supplemented medium for 1h, 4h, 22h, and 48h (+N). Three independent biological replicates were measured and the error bars represent the standard deviation.

Since nitrogen starvation limits consumption of carbon skeletons, starved *Synechocystis* 6803 cells store carbon polymers like glycogen. The stored glycogen content remained high in both strains during the first four hours after nitrogen addition (Fig. 3d). Thereafter, glycogen decreased in both strains but the degradation proceeded more rapidly in WT than in Δ*ssaA* (Fig. 3d). Our results indicate that although 6S RNA is not completely essential for the cells to recover from nitrogen starvation, the presence of 6S RNA accelerates the recovery process for hours.

Further investigations was carried out to analyze the 6S RNA transcript level under nitrogen starvation and recovery. Northern Blot analysis of 6S RNA revealed that the accumulation of 6S RNA increased in WT in the course of long term (189h -N) nitrogen depletion by ∼40 - 55% and returned to the original level, when normalized to 5S rRNA levels (Fig. 4a). However, since ribosomal RNA is rather expected to decrease during chlorosis [47] and consequently to increase again during recovery, 6S RNA accumulation is rather constant or slightly upregulated (Fig. 4a).

**Figure 4.**
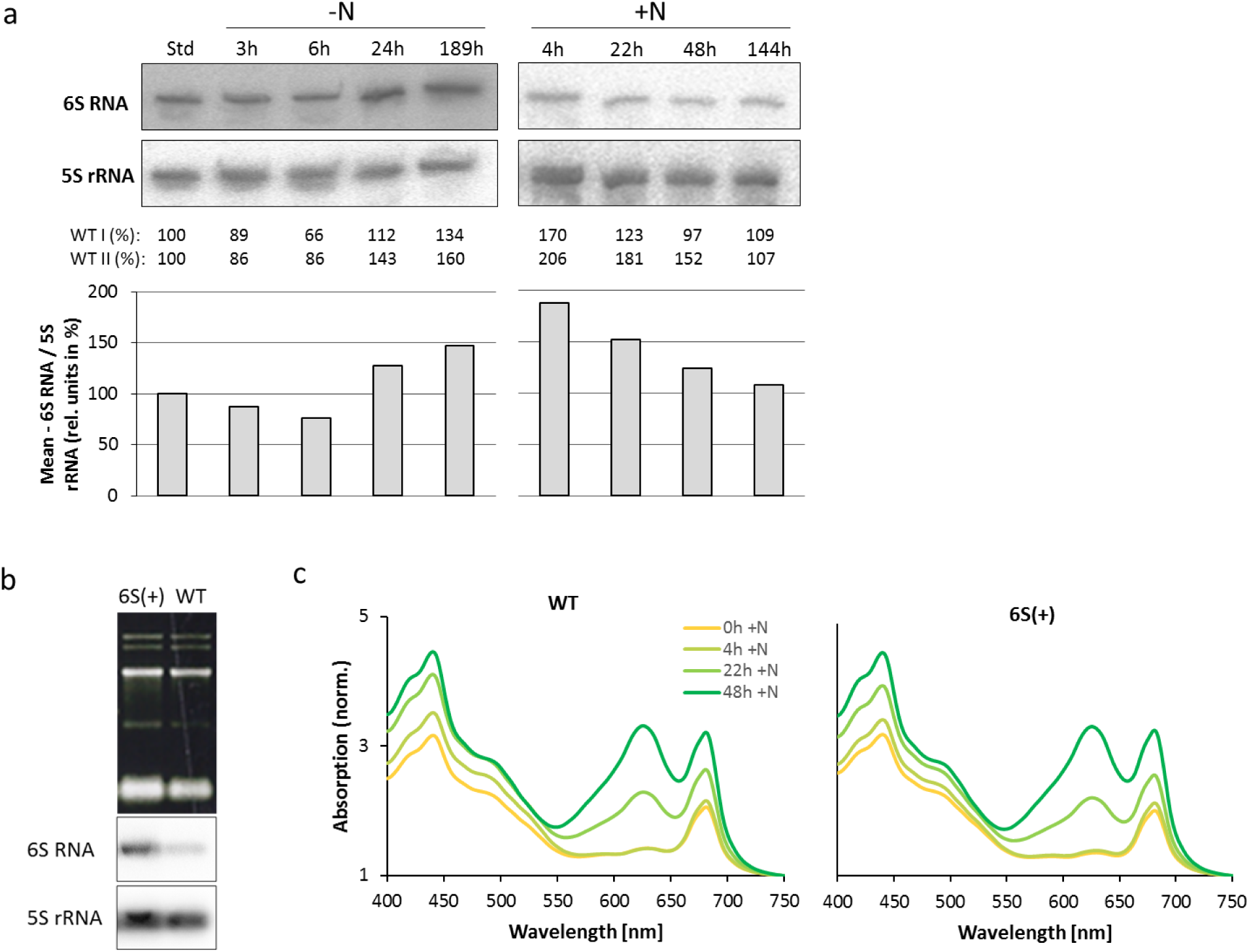
Analysis of 6S RNA transcript levels during nitrogen starvation and recovery in *Synechocystis* 6803 WT (a) and physiological characterization of the *Synechocystis* 6803 mutant strain overexpressing 6S RNA (6S(+)) (b and c). (a) Total RNA was isolated from cell cultures grown at standard growth conditions (t= Std), under nitrogen depleted conditions (t= 3h -N, t= 6h -N, t= 24h -N, t= 189h -N) and under nitrogen replete conditions (t= 4h +N, t= 22h +N, t= 48h +N, t= 144h +N) and samples with 5 μg RNA were analyzed by Northern blot hybridization. Autoradiograms are illustrated for WT I. 6S RNA and 5S rRNA signals of two biological replicates (WT I, WT II) were quantitated and 5S rRNA signal was used as an internal reference. The mean values are illustrated in the chart. Values of t= Std were set to 100 and samples of all other time points were calculated, respectively. (b) Northern Blot analysis to confirm overexpression of 6S RNA in 6S(+) mutant. 2 μg of total RNA was separated by electrophoresis on 1.3% agarose-formaldehyde gel. The amount of 6S RNA and 5S rRNA was detected by using radioactive [^32^P]-labeled oligonucleotides. (c) Physiological characterization of the 6S(+) strain. Absorbance spectra of the *Synechocystis* 6803 wild-type and the 6S(+) strain are illustrated for the time points t= 0h +N, t= 4h +N, t= 22h +N and t= 48h +N. The spectra were normalized at 750 nm.

Interestingly, overexpression of 6S RNA, as implemented in the 6S(+)-mutant, did not further accelerate recovery after nitrogen depletion (Fig. 4b and 4c).

### 6S RNA accelerates expression of household genes upon recovery from nitrogen starvation

Comparative transcriptome analysis was performed to infer a functional signature from the mutant-specific transcript profiles (raw data is available in supplementary files 2 and 3). For this purpose, microarray analysis was used to probe total RNA from WT and Δ*ssaA* cultures that were starved for nitrogen for seven days and recovered over a period of 22 hours. Sampling time points were t_1_ = 0h (7d -N; reference), t_2_=1h +N, t_3_=4h +N and t_4_=22h +N. At t_1_=0h a total set of 65 features showed differential accumulation (log2 fold change (FC) of > 1 (adjusted p-value of ≤ 0.05) in Δ*ssaA*, the majority of which were downregulated. Most of the latter comprised non-coding transcripts like asRNAs, but also eight mRNAs showed lower levels. Among them were several mRNAs encoding subunits of ATP synthase, the mRNA for cell division protein SepF as well as ferredoxin encoding *petF* and the high-light inducible *hliA*. The positively regulated genes included the heat stress responsive histidine kinase *hik34* and the chaperone encoding genes *dnaJ* and *groEL-2* (Fig. S3).

Temporally altered transcript levels in Δ*ssaA* from t_1_= 0h to all three analyzed time points of recovery were plotted as a function of the corresponding temporal changes in WT, distinctively comparing the recovery in both strains (Figs. S4-6). Comparison of transcript profiles indicated that only few genes were activated or repressed in the early phase of recovery, whereas 22h after addition of nitrogen, numerous genes responded, most of which were upregulated (Fig. S6).

Absolute expression values of selected groups of functionally related genes are further depicted in time courses in Fig. 5a-f. In WT cells transcriptional upshift of ATP synthase genes was seen already 1h after nitrogen addition and continued for whole recovery phase (Fig. 5a). In Δ*ssaA,* upregulation of *atp* transcripts was delayed, keeping *atp* transcripts at low levels for the first 4h.

**Figure 5.**
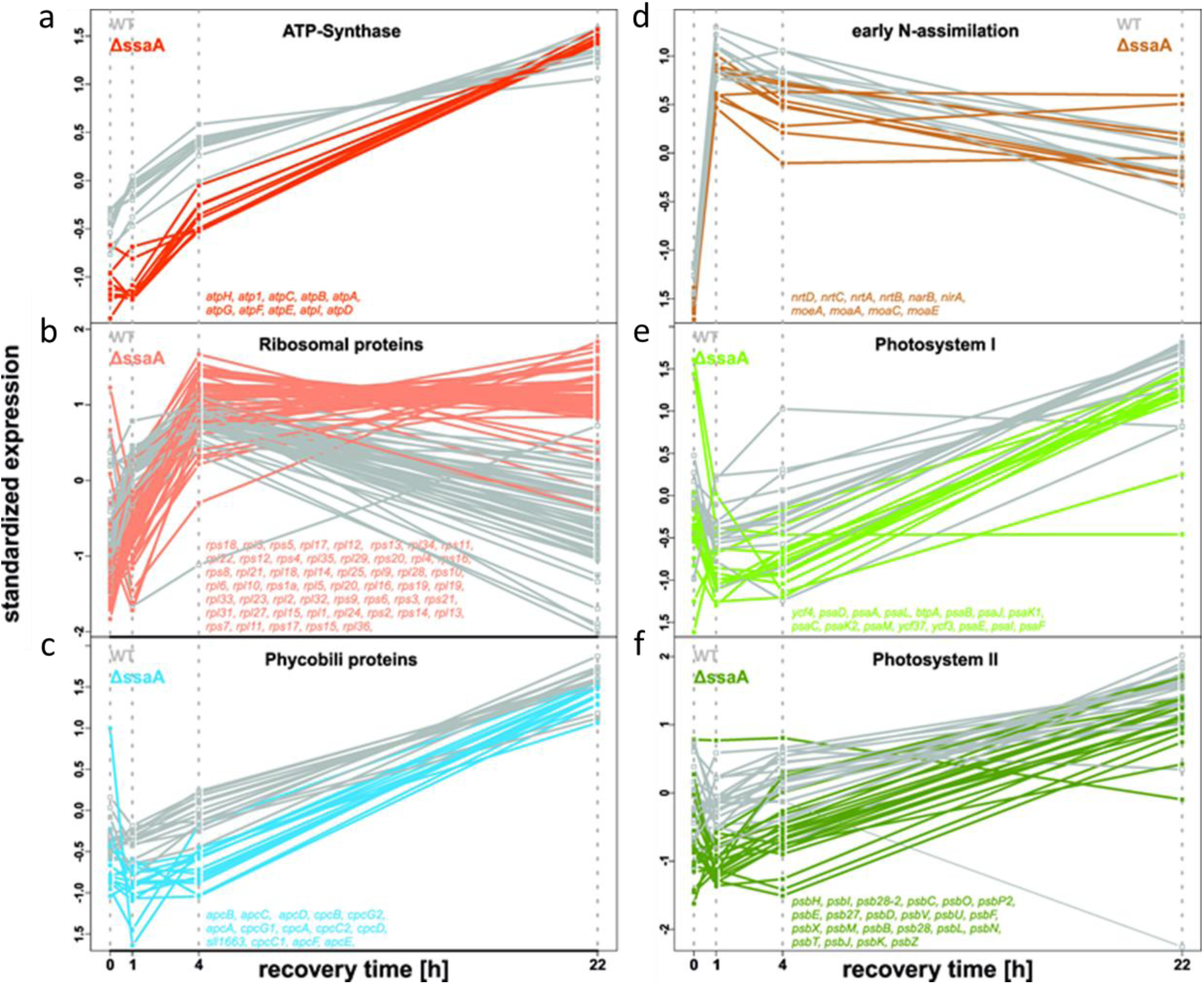
Temporal response of functional groups to nitrogen re-addition in the WT and Δ*ssaA*. Transcriptional profile of genes related to ATP synthase (a), ribosomal proteins (b), phycobili proteins (c), early nitrogen assimilation (d), PSI (e) and PSII (f) are grouped respectively. (x-axis) time after nitrogen addition. (y-axis) standardized normalized gene expression. To be able to compare the dynamics of the different genes in one plot the expression levels were standardized gene wise for both strains together, so that the average expression value is 0 and the standard deviation is one. WT expression is shown in grey and colored for Δ*ssaA* regarding to the color code in the scatter plots (Figs. S4-6). The genes which are plotted for each group are indicated in each sub-figure.

However, these differences between both strains vanished after 22h of recovery (Fig. 5a). Likewise, the transcriptional upshift of ribosomal subunits was delayed in Δ*ssaA* (Fig. 5b).

During the first 4h after nitrogen repletion many genes encoding proteins for the major photosynthetic complexes photosystem I (*psa* genes, Fig. 5e), photosystem II (*psb* genes, Fig. 5) and light harvesting phycobilisome antenna genes (Fig. 5f) produced less transcripts in Δ*ssaA* than in WT. t After 22h of recovery these effects wer not that clear anymore. For genomic context of *apc* and *cpc* operons, see Figure S7. Altogether, the transcriptomics data is in good agreement with the delayed physiological response of the mutant as reflected by the data shown in Figures 2 and 3.

Notably, levels of important ‘marker’ transcripts of nitrogen recovery encoding components of the nitrate uptake (*nrtABCD*) and early assimilation machinery (*narB*, *nirA*, *moaACE*) increased rapidly in both strains (Fig. 5d). Over the complete period, abundances of e.g. nitrate reductase encoding *nirA* kept stable in the mutant, whereas a moderate decline after the first hour was observed for all transcripts in the WT and likewise – for the majority of this group – in Δ*ssaA* (Fig. 5d).

For possible transcriptional regulators, we focused on genes encoding the regulatory σ factors. The *sigA* transcript (encoding primary σ factor SigA) was upregulated in Δ*ssaA* compared to WT after 4h of recovery, while transcripts for group 2 σ factors SigB and SigC were slightly up in all samples. Levels of *sigG*, encoding an ECF-type σ factor, were slightly down (Fig. 6a-g). Note that the mRNA accumulation of σ factor encoding genes in cannot considered a measure of σ factor recruitment by RNAP, but rather an effect of the latter.

**Figure 6.**
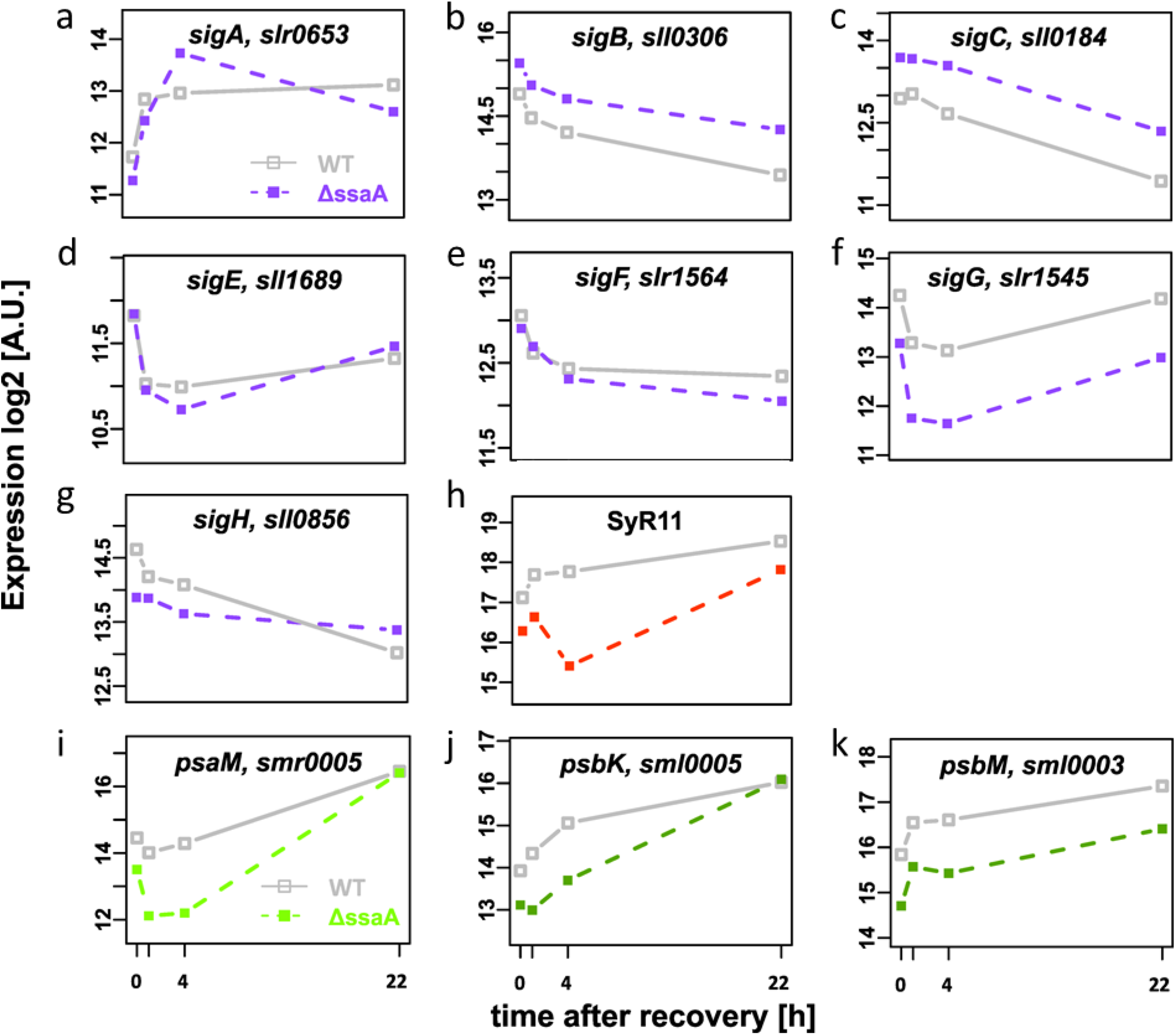
Temporal response of selected genes to nitrogen re-addition. Transcriptional profile of σ factors including group 1 σ factor SigA (a), group 2 σ factors SigB (b), SigC (c) and SigE (d) and group 3 σ factors SigF (e), SigG (f) and SigH (g) as well as of sRNA SyR11 (h). Representative mRNAs encoding photosystem genes (*psaM* (i); *psbK* (j) and *psbM* (k)) are shown to resolve the global effects illustrated in Fig. 5 exemplarily. (x-axis) time after nitrogen addition. (y-axis) normalized log2 expression. WT expression is shown in grey and Δ*ssaA* expression is color coded regarding to the color code in the scatter plots (Figs. S4-6).

In addition to protein-coding genes, a few small non-coding RNAs (sRNAs) showed altered levels in the mutant, including the previously characterized PmgR [48], NsiR4 [49] and PsrR1 [11], as well as several hypothetical riboregulators that were identified by differential RNA sequencing (dRNA-seq) [50]. Among these transcripts, SyR11 exhibited the most diverging accumulation kinetics (Fig. 6h; see also Northern Blot validation in Figure S8). Additionally, Fig. 6i-k illustrate the mRNA abundances of three exemplarily selected transcripts (*psaM, psbK, psbM*) of PSI and PSII, revealing the decelerated upregulation of components of the photosynthetic apparatus in Δ*ssaA* in detail.

### 6S RNA determines the allocation of SigA, SigB, SigC and SigE in the RNA polymerase holoenzyme

We next studied if the absence of 6S RNA influences the content of the primary or group 2 σ factors and/or recruitment of different σ factors by the RNAP core to form the holoenzyme during nitrogen starvation and recovery phase. To that end, the γ-subunit of RNAP core was replaced with His-tagged γ-subunit in WT and δ*ssaA* as described recently for a non-motile *Synechocystis* 6803 strain [51]. The WT RNAP-His and Δ*ssaA* RNAP-His strains were grown under standard conditions or subjected to nitrogen depletion for 6 days; recovery samples were taken 1h, 4h or 22h after nitrogen addition.

The His-tagged RNAP complexes were collected from one-half of each sample and the soluble proteins were isolated from the other half.

To measure possible changes in SigA content and/or in recruitment efficiencies by the RNAP core, isolated soluble proteins and collected RNAP complexes were separate on SDS-PAGE, respectively. Subsequently, the SigA content was detected using a SigA specific antibody [51]. Then membranes were re-probed with the antibody against the α (or β) subunit of the RNAP core and the SigA content was normalized to the RNAP core content. Finally, after each treatment, the amount of SigA was compared to the amount of SigA in cultures from standard (nitrogen replete) conditions.

SigA almost vanished from the RNAP holoenzyme during nitrogen depletion in both strains (Fig. 7a), but SigA protein content decreased by only 65% (Fig. 7b) indicating that the recruitment of SigA by RNAP core reduced more drastically than SigA content. After nitrogen addition, RNAP core successively recruited more SigA (Fig. 7a). The recovery of the SigA recruitment occurred more slowly in Δ*ssaA* than in WT (Fig. 7a), although the SigA content increased more rapidly during recovery in Δ*ssaA* than in WT (Fig. 7b). Poor recruitment of SigA in Δ*ssaA* is an obvious reason for slow activation of household genes in this strain.

**Figure 7.**
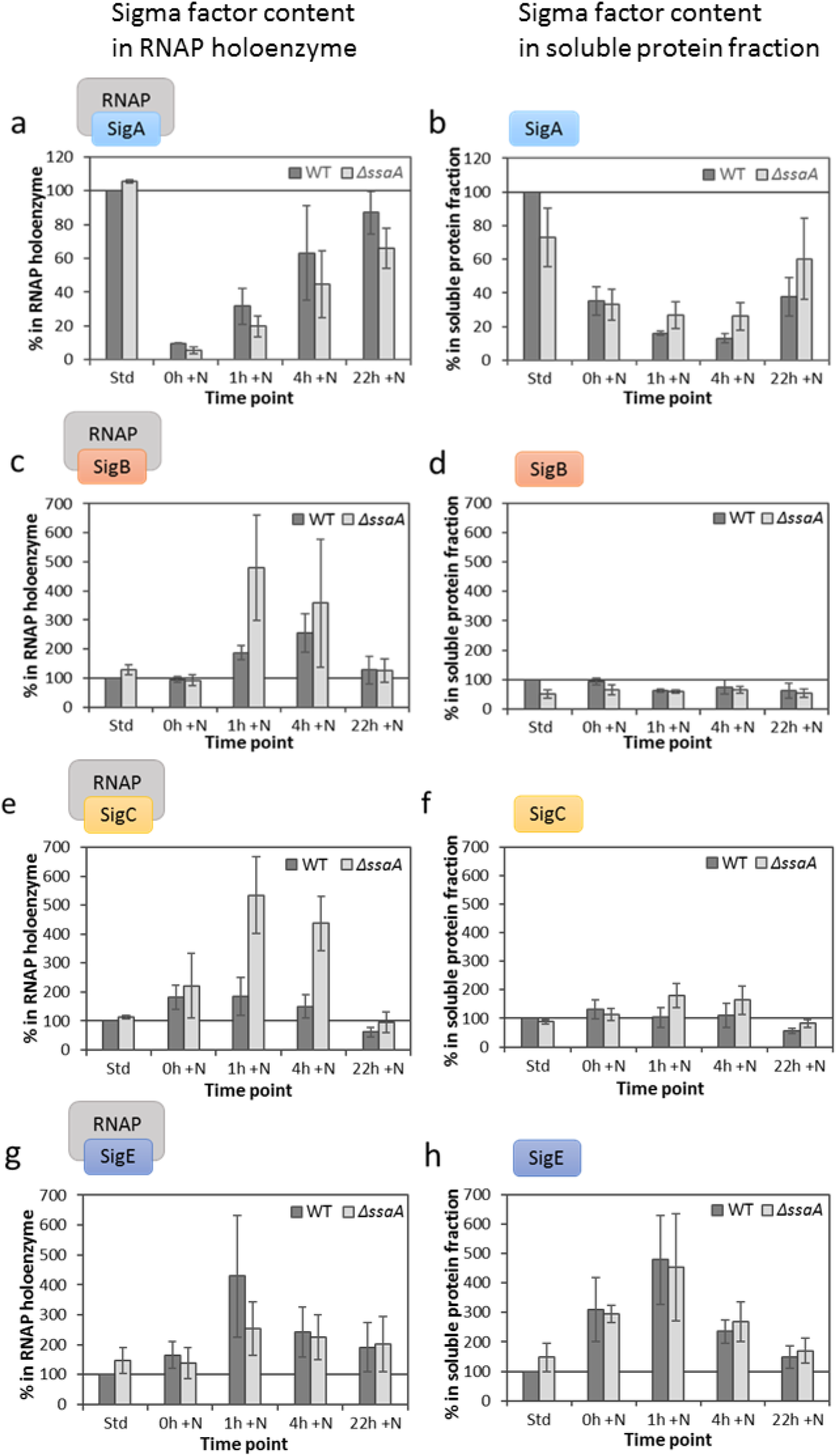
Relative content of different sigma factors (σ) in RNAP holoenzyme (left panel; a, c, e, g) and in the soluble protein fraction (right panel; b, d, f, h) of WT-RNAP-HIS (WT) and Δ*ssaA*-RNAP-HIS mutant (Δ*ssaA*) strain under standard conditions (Std), under nitrogen deficiency for 6 days (t= 0h +N) and during recovery (t= 1h +N, t= 4h +N and t= 22h +N). Equal amounts of collected RNAP complexes or soluble proteins were separated on SDS-PAGE, group 1 σ factor SigA (a, b) and group 2 σ factors SigB (c, d), SigC (e, f) and SigE (g, h) were detected by immunoblot analysis. The σ factor content was normalized to α or β RNAP core subunit. For each σ factor the value of the WT-RNAP-HIS strain at standard conditions (Std) was set to 100 and the other samples were compared respectively. Three independent biological replicates were analyzed and the error bars represent the standard error.

When group 2 σ factors were analyzed, using the same method as described for the primary σ factor, the results showed that in nitrogen starved WT cells (0h +N) both recruited and total SigB remained at the same level as under standard conditions (Fig. 7c and 7d). Although the total amount of SigB decreased moderately during the recovery period, this σ factor was recruited in particular during the early recovery phase (Fig. 7c and 7d). In Δ*ssaA,* SigB was more efficiently recruited by RNAP core both under standard conditions and during early recovery than in WT (Fig. 7c and 7d).

Thus, 6S RNA seems to restrict recruitment of SigB by the RNAP core. Like SigB, SigC was highly recruited during recovery phase, however SigC was already recruited efficiently in nitrogen depleted cells (Fig. 7e and 7f). Very high levels of SigC recruitment were observed in the mutant strain at early time points of recovery. The amount of SigD was very low or below detection limit in all samples and quantification of SigD results was not possible. Unlike the other σ factors, the SigE protein content was clearly lower in nitrogen depleted samples, which was also observed in samples from the early recovery phase. Increased SigE content resulted in enhanced recruitment of SigE during nitrogenrecovery, especially in early phase (Fig. 7g and 7h).

## Discussion

### 6S RNA promotes the recovery process from N starvation

The acclimation processes of cyanobacteria to changing nitrogen availability are underlain by a complex regulatory network, involving signal-driven transcriptional regulons and differential σ factor activity, refined with a layer of sRNA-mediated post-transcriptional regulation [49, 52-54].

Here we demonstrate that the cyanobacterial 6S RNA homologue plays a role within the rapid transcriptional response during recovery from nitrogen depletion. The Δ*ssaA* mutant of *Synechocystis* 6803 showed a more sustained bleaching phenotype linked with a generally decelerated transcriptional and physiological response during recovery. Thus, within this gradual acclimation process 6S RNA containing cells have a substantial selective advantage over cells missing 6S RNA. On the other hand, overproduction of 6S RNA in *Synechocystis* did not accelerate recovery from nitrogen starvation, suggesting that this riboregulator is not a limiting factor in the native acclimation system.

The behavior of 6S RNA during nitrogen-deficiency-induced bleaching and subsequent recovery in *Synechocystis* 6803 differs from that reported for stationary phase in heterotrophic bacteria. In *Synechocystis* 6803, 6S RNA increases only slightly during nitrogen starvation (Fig. 4a) and 6S RNA is quite abundant under standard conditions [50, 55]. For the cyanobacterium *Synechococcus* sp. PCC 6301 it has been reported previously, that 6S RNA levels decrease during stationary phase [27]. In *E. coli,* 6S RNA molecules enrich 10-fold from early exponential growth phase to late stationary phase [19] and binding of RNAP-σ^70^ holoenzyme to 6S RNA keeps the RNAP-σ^70^ holoenzyme complex abundant although it becomes inactive. The importance of the regulatory function of 6S RNA in *E. coli* is obvious as deletion of 6S RNA affects expression of ∼ 800 genes in stationary phase [56]. Contrary to that, 6S RNA might play only a minor role in nitrogen starved *Synechocystis* 6803 cells as only very few genes show different expression in nitrogen starved WT and Δ*ssaA* cells. Furthermore, the amount of the RNAP-SigA complex drastically reduces during nitrogen starvation in *Synechocystis* 6803 indicating that formation of inactive RNAP-SigA-6S RNA holoenzyme complexes is not the main regulatory mechanism behind reduced transcription of household genes. It is possible that 6S RNA exhibits some functional diversity within different bacterial phyla.

Interestingly, *Bacillus subtilis* produces two different 6S RNAs, 6S-1 RNA and 6S-2 RNA [57]. The 6S-1 RNA (*bsrA*) is highly upregulated in stationary phase just like 6S RNA in *E. coli* while 6S-2 RNA (*bsrB*) transcript level decrease towards stationary phase [58]. Therefore, *Synechocystis* 6803 6S RNA possibly resembles 6S-2 RNA in matters of transcriptional regulation.

In accordance with the physiological data, transcriptome profiling demonstrated that particularly genes encoding ATP synthase, PSI and PSII and phycobilisomes as well as the translational machinery were overall negatively affected by the absence of 6S RNA during recovery from nitrogen starvation. The resuscitation process from severe long-term nitrogen starvation has recently been described in detail for *Synechocystis* 6803 by Klotz and co-workers [45]. The recovery process from nitrogen starvation can be divided into two phases. Phase 1 is characterized by regeneration of basic cellular functions, including respiration, translation and nitrogen assimilation, which could be clearly confirmed by our transcriptome study. Here, certainly expression of ATP synthase and ribosomal genes showed a delay of general upshift from 1h to 4h after nitrogen addition in the Δ*ssaA* mutant. The kinetics of photosynthesis-related mRNA accumulation further emphasizes the generally delayed recovery: the early phase 2 response, which is partially initiated after 4h in WT, rather arises later in the mutant, largely approaching WT levels after 22h. Moreover, due to delayed transcriptional activation of phase 1 and phase 2 genes, recovery of physiological processes like phycobilisome reassembly, photosynthetic activity and glycogen degradation were delayed in Δ*ssaA*.

Our results show that 6S RNA regulates recruitment of σ factors especially during early recovery from nitrogen depletion but does not affect expression of NtcA controlled genes like *amt1*, *glnA*, *glnB* or *sigE*, indicating an additional regulatory circuit that acts independently and putatively supplementary to the well-characterized PipX-PII-NtcA network [46, 47]. Although none of the group 2 σ factors alone is essential for acclimation to nitrogen deficiency and subsequent recovery, cells without functional group 2 σ factors were more sensitive to nitrogen depletion [30].

During recovery from nitrogen depletion WT cells recruited SigA more efficiently than Δ*ssaA* cells, indicating that SigA recruitment was promoted by 6S RNA when cells switched from nitrogen depletion to repletion.

The amount of RNAP-SigC increased in both strains during nitrogen deficiency and in the beginning of recovery phase. However, the SigC recruitment during the early recovery phase was considerably more pronounced in Δ*ssaA* than in WT. Cells containing SigC as the only functional group 2 σ factor (the Δ*sigBDE* strain) showed a delayed recovery from nitrogen starvation [42] and SigC has been suggested to regulate nitrogen metabolism in stationary phase [59]. Altogether, these results suggest that recruitment of SigC by RNAP core prevent exit of cells from the stress-adapted state.

Likewise, SigB was recruited in larger quantities in Δ*ssaA* at the beginning of recovery. Cells without stress inducible SigB are vulnerable to many stress conditions including heat [60] and high salt [61, 62], and SigB helps to keep photosynthesis active under nitrogen deficiency [42]. It remains to be elucidated which role SigB might play in activation of the *sigA* gene during recovery from nitrogen deficiency induced stationary phase. However, a possible connection can be derived from previous experiments with dark/ light transitions [63].

The amount of SigE increases during nitrogen starvation and early recovery in both strains (Fig. 7h). However, the recruitment of SigE was less efficient in Δ*ssaA* than in WT in the early recovery phase. SigE has been earlier suggested to function in nitrogen stress responses [38, 41], and its role in triggering sugar catabolism is well-characterized [41, 43, 64]. Since *glgX* and *glgP* are positively controlled by SigE [65], the observed effect on SigE recruitment might underlie the decelerated glycogen degradation of the mutant strain. An anti-σ factor, ChlH, has been proposed to inhibit SigE recruitment in a light-dependent manner [66]. Moreover, the sRNA SyR11, which is presumably involved in post-transcriptional activation of *sigE* expression (unpublished results), showed significantly reduced levels in Δ*ssaA*, particularly during early recovery. Therefore, more detailed research on the involvement of integrated post-translational effects and - certainly - riboregulatory events will further disclose the complexity of the cyanobacterial σ factor network during physiological transitions. While not a conspicuous feature in this comparative study, the asRNA slr0653-as4 appears to be a potent inhibitor of *sigA* expression during starvation [45], exemplifying the significance of the RNA-regulatory layer in this process.

## Conclusion

Summarizing our current knowledge, we suggest a probable mechanism for 6S RNA-dependent regulation within the acclimation process to changing nitrogen supply. In *Synechocystis* 6803, formation of RNAP-SigA holoenzymes drastically decreases during nitrogen deprivation while recruitment of different group 2 σ factors either increases (SigC and SigE) or remains at the same level (SigB and SigD) as under standard conditions. After prolonged nitrogen deficiency, transcription activity is low in general [45], most probably because expression of household genes is depressed due to low formation of RNAP-SigA holoenzymes. During the recovery phase, group 2 σ factors are first to be efficiently recruited by RNAP core, in order to be replaced later on by the primary σ factor. 6S RNA regulates this process by facilitating the replacement of group 2 σ factors by the primary σ factor. Due to a delayed formation of RNAP-SigA holoenzymes in the Δ*ssaA* strain, activation of many household genes occurs at a slower rate resulting in a decelerated re-synthesis and activation of light harvesting and photosynthetic complexes.

Hence, with our study we extend the established function of 6S RNA towards the recruitment of housekeeping-versus-stress σ factors by RNAP to cyanobacteria. Thereby, 6S RNA is mediating their transcriptional response and supports a rapid adaptation to nutrient - in this case nitrogen - depletion and repletion. Further, our findings imply that cyanobacterial 6S RNA might discriminate not only between group 1 and group 2 but also between the different group 2 σ factors.

## Methods

### Bacterial strains, growth conditions and experimental setup

The *Synechocystis* sp PCC 6803 strain PCC-M [67] used in this study was provided by S. Shestakov (Moscow State University) and cultivated on 1% (w/v) agar (Bacto agar; Difco) plates containing BG- 11 mineral medium [68]. Liquid cultures of WTand mutant strains were grown in BG-11 medium containing 10 mM TES buffer, pH 8.0, under continuous illumination with white light at 80 µmol photons m–^2^ s–^1^ at 30 °C, supplemented with a continuous stream of ambient air. BG11 plates for the cultivation of mutant strains were supplemented with 40 μg ml^-1^ kanamycin (Δ*ssaA*) and 10 μg ml^-1^ chloramphenicol (WT-RNAP-HIS) or both antibiotics (Δ*ssaA*-RNAP-HIS), respectively. No antibiotics were added to the liquid cultures. Deviating growth conditions are specified in the corresponding figure legends or in the text. For nitrogen depletion, NaNO_3_ was omitted from the medium and cells were first washed twice and then resuspended in nitrogen-free medium. Cultures were incubated in nitrogen-depleted medium for 3-8 days. To induce nitrogen recovery, starved cells were supplemented with 17.6 mM NaNO_3_. Cells were harvested after cultivation in nitrogen-repleted medium for the indicated times (1h, 4h, 22h, 48h, 144h).

### Construction of 6S RNA mutants and RNAP-HIS mutants

For the construction of a 6S RNA deletion mutant (Δ*ssaA*), the complete *Synechocystis* 6803 sequence region of the *ssaA* gene was deleted by replacement with a kanamycin-resistance cassette (Km^R^), which was inserted between the chromosomal nucleotide positions 1886182 and 1887765. As distinguished from the previously described Δ*ssaA* mutant [28], the Km^R^ cassette harbored its own promotor and was flanked by ∼700 bp regions upstream and downstream of the *ssaA* gene for homologous recombination. The Δ*ssaA-*mutant was selected on BG11 agar plates containing 50 μg ml^-1^ kanamycin. PCR analysis was used to verify complete segregation of WT chromosomal copies (Fig. 1B). The primers P1 (Δ*ssaA*_700up-Fw) and P2 (Δ*ssaA*_700down-Rv) are listed below in Table S1. To create a strain overexpressing 6S RNA (6S(+)), the *ssaA* gene locus and its promoter region (∼150 bp of upstream region) were inserted into the self-replicating vector pVZ321 [69], containing a chloramphenicol resistence gene using XhoI and XbaI. The vector was then transferred to WT cells by conjugation. Exconjugants were selected on BG11 agar plates containing 10 µg ml^-1^ chloramphenicol. Complementation of the Δ*ssaA* strain (Δ*ssaA*-c) was achieved by transferring the pVZ321 vector containing the *ssaA* gene plus ∼150 bp of its upstream region and a chloramphenicol resistance gene to Δ*ssaA* mutant cells by conjugation. The Δ*ssaA*-c-mutant was selected on BG11 agar plates containing 50 μg ml^-1^ kanamycin and 10 µg ml^-1^ chloramphenicol. For the generation of a WT-RNAP-HIS and Δ*ssaA*-RNAP-HIS mutant strain, *Synechocystis* 6803 WT and Δ*ssaA* cells were transformed using the plasmid pC-HIS-rpoC1-CAP-I-I (containing a nine amino acids long His-tag attached to the C terminal end of the Δ subunit, and a chloramphenicol resistance cassette after the *rpoC1* gene) which was described earlier by Koskinen *et al*. [51]. WT-RNAP-HIS and Δ*ssaA*-RNAP-HIS mutants were probed on BG11 plates containing 10 µg ml^-1^ chloramphenicol and full segregation of the mutants was tested by PCR analysis using primers rpoC1His5 and rpoC1His6 (Table S1) from Koskinen *et al*. [51].

### Analysis of growth, pigmentation and photosynthetic capacity

Whole cell absorbance spectra were recorded from 550 to 750 nm on a Specord(r) 200 PLUS spectrophotometer (Analytik Jena, Jena, Germany) and were corrected for a residual scattering at 750 nm. Chlorophyll was extracted in 90% methanol and the content was measured by spectrophotometry [70]. Contents of phycocyanin and allophycocyanin were determined in the soluble protein fraction according to Tandeau de Marsac and Houmard [70]. The pigment contents were normalized to the optical density at 750 nm. Light saturated oxygen evolution rates of the WT and Δ*ssaA* mutant strain were measured with a Clark-type oxygen electrode (Hansatech) in BG-11 medium *in vivo* in the presence of 10 mM NaHCO_3_ at 32 °C, illuminated with white light (photosynthetic photon flux density 3000 μmol m^-2^ s^-1^) [61]. To determine PSII activity samples were supplemented with 1 mM 2,6-dimethoxybenzoquinone (DMBQ) and 1 mM ferricyanide. In preparation for 77 K fluorescence emission spectroscopy, cells were concentrated to 35 μg Chl ml ^-1^, and 50 μl samples were immediately frozen in liquid nitrogen. The fluorescence was measured at 77 K with orange light excitation (580 nm) using an Ocean Optics S2000 spectrometer. The spectra were corrected with a moving median, subtraction of the dark level and division by the peak value of PSI at 721 nm and setting this value to 1.

### Determination of glycogen content

The content of glycogen per ml cell suspension was determined as described by Gründel *et al*. [71]. Cell pellets of 0.5 - 1.5 ml culture were suspended in 30% (w/v) KOH and incubated for 2 h at 95 °C.

An addition of ice-cold ethanol to a final concentration of 75% (v/v) and a following incubation on ice for at least 1.5 h were required for the precipitation of glycogen. The isolated glycogen was centrifuged for 15 min (16000 *g*, 4 °C) and the pellet was washed with 70% and 98% (v/v) ice-cold ethanol and dried for 10 min at 60 °C. For the enzymatic hydrolysis of glycogen to glucose, the pellet was suspended in 100 mM sodium acetate (pH 4.5) including 2 mg/ml amyloglycosidase and incubated for 4 h at 95 °C. To determine the content of glucose, a hexokinase reagent was used according to the manufacturer´s protocol and measured spectrophotometrically.

### Isolation of RNA polymerase complex and immunoblot analysis

Cells from WT-RNAP-HIS and Δ*ssaA*-RNAP-HIS mutant strain (50 ml, OD_750nm_ = 0.8 - 1.1) were collected from growth at standard conditions, after 6d of growth in nitrogen depleted medium or after 1h, 4h and 22h of recovery in nitrogen replete medium. The samples were broken and the soluble protein fraction was obtained as described in Koskinen *et al*. [51]. For the isolation of RNAP the Dynabeads(r)HIS-Tag Isolation and Pulldown Kit was applied according to the manufacturer´s instructions. 900 μg of soluble protein fraction was used for the pulldown and RNAP complexes were eluted to 75 μl of elution buffer [51]. 10 μl of His-tag purified RNAP samples were subjected to 10% NEXT GEL SDS-PAGE according to the protocol of Koskinen *et al*. [51]. RNAP complexes were transferred to Immobilon-P membrane using the semidry blotting method followed by incubation with specific polyclonal antibodies against σ factors SigA, SigB, SigC and SigE [51, 72, 73]. Proteins were visualized and quantified as described by Koskinen *et al*. [51]. Membranes were re-probed with antibodies against α or β subunits of the RNAP core [73] and the σ factor content was normalized to α (SigA, SigC, SigE) or β subunit content (SigB). Similarly, 5 μg of soluble proteins for SigA, 25 μg for SigB, 15 μg for SigC and 25 μg for SigE were used for separation on SDS-PAGE and were detected as described above.

### RNA isolation and Northern blot hybridization

For the isolation of RNA, samples from discrete stages of cultivation were taken at the indicated times and were immediately put on ice and spun down at 4 °C. The preparation of total RNA from *Synechocystis* 6803 was essentially performed as described previously [74], using the Hot Trizol method, followed by 2-propanol precipitation. To determine the concentration and to check the quality of total RNA, we made use of a NanoDrop N-10000 spectrophotometer and electrophoretic separation on 1.3% agarose-formaldehyde gels. The correct deletion of the *ssaA* gene was tested by northern blot analysis using a directed [^32^P]-labelled DNA oligonucleotide for 6S RNA. Additionally, directed [^32^P]-labelled DNA oligonucleotides for 6S RNA and 5S rRNA were used to analyze the 6S RNA level. For the detection of SyR11 transcript levels RNA samples (2 μg) were separated on 10%- urea-PAA gels and transferred to Hybond^TM^-N^+^ membrane. The RNA was hybridized against single-stranded DIG-labelled RNA oligonucleotides overnight at 62 °C using the DIG Northern Starter Kit.

DIG-labelled RNA oligonucleotides were prepared according to the manufacturer´s instructions using *in vitro* transcription with the AmpliScribe kit (T7-Flash Transcription kit, Epicentre) from a PCR product containing a T7 promotor. The PCR fragment was amplified with the primer pair SyR11_fw and SyR11_T7_rev (Table S1).

### Microarray analysis

The microarray design, hybridization procedure and data analysis have been described previously [50, 75]. For the microarray analysis total RNA samples from two biological replicates were used. Raw data were processed with the R package limma. Median signal intensity was cyclic loess normalized. The p values were generated by limma [76] and adjusted for multiple testing with the Benjamini Hochberg method.

## Abbreviations

σ, sigma factor; *B. subtilis*, *Bacillus subtilis;* Chl, chlorophyll *a*; Cm^r^, chloramphenicol resistance cassette; DMBQ, 2,6-dimethoxybenzoquinone; dRNA-seq, differential RNA sequencing; DIG, digoxygenin; *E. coli, Escherichia coli;* Km^r^, kanamycin resistance cassette; N; nitrogen; ncRNA, non-coding RNA; nt, nucleotide; OD, optical density; PBS, phycobilisome; pRNA, product RNA; PSI, photosystem I; PSII, photosystem II; RNAP, RNA polymerase; sRNA, small RNA; std, standard; *Synechocystis* 6803, *Synechocystis* sp. PCC 6803; WT, wild type

## Declarations

## Acknowledgements

We thank Viktoria Reimann for technical support and Anne Rediger for generating the Δ*ssaA* mutant and the 6S (+) mutant strain.

## Authors´ contributions

I.M.A. and D.D. designed and supervised the project. B.H. carried out all physiological studies, analyzed photosynthetic parameters, prepared RNA for microarray analysis, contributed to the recruitment analysis and performed Northern Blot hybridization. J.G. and W.R.H. performed the microarray analysis, K.H. contributed to the recruitment and soluble protein analysis. B.H. and D.D. conducted all verification experiments. I.M.A., D.D., B.H., J.G. and T.T. interpreted the data. B.H. and J.G. prepared the figures. B.H., D.D., J.G., W.R.H., T.T., K.H. and I.M.A. drafted the manuscript. All authors reviewed and approved the manuscript.

## Availability of data and materials

The microarray data generated during the current study are available in the database [GEO: <u>GSE83387</u>].

## Consent for publication

Not applicable.

## Ethics approval and consent to participate

Not applicable.

## References

[1] Möllers KB, Cannella D, Jørgensen H, Frigaard N-U. Cyanobacterial biomass as carbohydrate and nutrient feedstock for bioethanol production by yeast fermentation. Biotechnology for Biofuels. 2014;7:64-.

[2] Dexter J, Armshaw P, Sheahan C, Pembroke JT. The state of autotrophic ethanol production in Cyanobacteria. Journal of Applied Microbiology. 2015;119:11–24.

[3] Sarsekeyeva F, Zayadan BK, Usserbaeva A, Bedbenov VS, Sinetova MA, Los DA. Cyanofuels: biofuels from cyanobacteria. Reality and perspectives. Photosynth Res 2015;125:329–40.

[4] Erdrich P, Knoop H, Steuer R, Klamt S. Cyanobacterial biofuels: new insights and strain design strategies revealed by computational modeling. Microbial Cell Factories. 2014;13:128.

[5] Rögner M. Metabolic engineering of cyanobacteria for the production of hydrogen from water. Biochemical Society Transactions. 2013;41:1254–9.

[6] Angermayr SA, van der Woude AD, Correddu D, Vreugdenhil A, Verrone V, Hellingwerf KJ. Exploring metabolic engineering design principles for the photosynthetic production of lactic acid by Synechocystis sp. PCC6803. Biotechnology for Biofuels. 2014;7:99-.

[7] Pattanaik B, Lindberg P. Terpenoids and Their Biosynthesis in Cyanobacteria. Life. 2015;5:269–93.

[8] Ducat DC, Way JC, Silver PA. Engineering cyanobacteria to generate high-value products. Trends in Biotechnology. 2011;29:95–103.

[9] Duhring U, Axmann IM, Hess WR, Wilde A. An internal antisense RNA regulates expression of the photosynthesis gene isiA. Proc Natl Acad Sci U S A. 2006;103:7054–8.

[10] Hernández-Prieto MA, Schön V, Georg J, Barreira L, Varela J, Hess WR, et al. Iron Deprivation in Synechocystis: Inference of Pathways, Non-coding RNAs, and Regulatory Elements from Comprehensive Expression Profiling. G3: Genes|Genomes|Genetics. 2012;2:1475–95.

[11] Georg J, Dienst D, Schurgers N, Wallner T, Kopp D, Stazic D, et al. The small regulatory RNA SyR1/PsrR1 controls photosynthetic functions in cyanobacteria. Plant Cell. 2014;26:3661–79.

[12] Eisenhut M, Georg J, Klähn S, Sakurai I, Mustila H, Zhang P, et al. The Antisense RNA As1_flv4 in the Cyanobacterium Synechocystis sp. PCC 6803 Prevents Premature Expression of the flv 4–2 Operon upon Shift in Inorganic Carbon Supply. The Journal of Biological Chemistry. 2012;287:33153–62.

[13] Gierga G, Voss B, Hess WR. Non-coding RNAs in marine Synechococcus and their regulation under environmentally relevant stress conditions. The ISME Journal. 2012;6:1544–57.

[14] Klähn S, Schaal C, Georg J, Baumgartner D, Knippen G, Hagemann M, et al. The sRNA NsiR4 is involved in nitrogen assimilation control in cyanobacteria by targeting glutamine synthetase inactivating factor IF7. Proceedings of the National Academy of Sciences of the United States of America. 2015;112:E6243–E52.

[15] Barrick JE, Sudarsan N, Weinberg Z, Ruzzo WL, Breaker RR. 6S RNA is a widespread regulator of eubacterial RNA polymerase that resembles an open promoter. RNA. 2005;11:774–84.

[16] Steuten B, Hoch PG, Damm K, Schneider S, Kohler K, Wagner R, et al. Regulation of transcription by 6S RNAs: insights from the Escherichia coli and Bacillus subtilis model systems. RNA Biol. 2014;11:508–21.

[17] Cavanagh AT, Wassarman KM. 6S RNA, a Global Regulator of Transcription in Escherichia coli, Bacillus subtilis, and Beyond. Annual Review of Microbiology. 2014;68:45–60.

[18] Burenina OY, Elkina DA, Hartmann RK, Oretskaya TS, Kubareva EA. Small noncoding 6S RNAs of bacteria. Biochemistry (Moscow). 2015;80:1429–46.

[19] Wassarman KM, Storz G. 6S RNA Regulates E. coli RNA Polymerase Activity. Cell. 2000;101:613–23.

[20] Trotochaud AE, Wassarman KM. 6S RNA function enhances long-term cell survival. J Bacteriol. 2004;186:4978–85.

[21] Cavanagh AT, Klocko, A.D., Liu, X., and Wassarman, K.M. Promoter specificity for 6S RNA regulation of transcription is determined by core promoter sequences and competition for region 4.2 of sigma(70). Mol Microbiol. 2008;67:1242–56.

[22] Neusser TP, T.; Geissen, R.; Wagner, R. Depletion of the non-coding regulatory 6S RNA in E. coli causes a surprising reduction in the expression of the translation machinery. BMC Genomics. 2010;11:165.

[23] Wassarman KM, Saecker RM. Synthesis-Mediated Release of a Small RNA Inhibitor of RNA Polymerase. Science. 2006;314:1601–3.

[24] Gildehaus N, Neußer T, Wurm R, Wagner R. Studies on the function of the riboregulator 6S RNA from E. coli: RNA polymerase binding, inhibition of in vitro transcription and synthesis of RNA-directed de novo transcripts. Nucleic Acids Research. 2007;35:1885–96.

[25] Wurm R, Neußer T, Wagner R. 6S RNA-dependent inhibition of RNA polymerase is released by RNA-dependent synthesis of small de novo products. Biol Chem. 2010;391:187–96.

[26] Axmann IM, Kensche P, Vogel J, Kohl S, Herzel H, Hess WR. Identification of cyanobacterial non-coding RNAs by comparative genome analysis. Genome Biol. 2005;6:R73.

[27] Watanabe T, Sugiura M, Sugita M. A novel small stable RNA, 6Sa RNA, from the cyanobacterium Synechococcus sp. strain PCC6301. FEBS Letters. 1997;416:302–6.

[28] Rediger A, Geissen R, Steuten B, Heilmann B, Wagner R, Axmann IM. 6S RNA - an old issue became blue-green. Microbiology. 2012;158:2480–91.

[29] Trotochaud AE, Wassarman KM. 6S RNA Regulation of pspF Transcription Leads to Altered Cell Survival at High pH. Journal of Bacteriology. 2006;188:3936–43.

[30] Hoch PG, Burenina OY, Weber MHW, Elkina DA, Nesterchuk MV, Sergiev PV, et al. Phenotypic characterization and complementation analysis of Bacillus subtilis 6S RNA single and double deletion mutants. Biochimie. 2015;117:87–99.

[31] Cavanagh AT SJ, Wassarman KM. Regulation of 6S RNA by pRNA synthesis is required for efficient recovery from stationary phase in E. coli and B. subtilis. Nucleic Acids Res. 2012;40:2234–46.

[32] Cabrera-Ostertag IJ, Cavanagh AT, Wassarman KM. Initiating nucleotide identity determines efficiency of RNA synthesis from 6S RNA templates in Bacillus subtilis but not Escherichia coli. Nucleic Acids Research. 2013;41:7501–11.

[33] Trotochaud AE, Wassarman KM. 6S RNA function enhances long-term cell survival. J Bacteriol. 2004;186.

[34] Cavanagh AT, Klocko AD, Liu X, Wassarman KM. Promoter specificity for 6S RNA regulation of transcription is determined by core promoter sequences and competition for region 4.2 of sigma(70). Mol Microbiol. 2008;67.

[35] Allen MM, Smith AJ. Nitrogen chlorosis in blue-green algae. Arch Mikrobiol. 1969;69:114–20.

[36] Collier JL, Grossman AR. Chlorosis induced by nutrient deprivation in Synechococcus sp. strain PCC 7942: not all bleaching is the same. J Bacteriol. 1992;174:4718–26.

[37] Görl M, Sauer J, Baier T, Forchhammer K. Nitrogen-starvation-induced chlorosis in Synechococcus PCC 7942: adaptation to long-term survival. Microbiology. 1998;144 (Pt 9):2449–58.

[38] Krasikov V, Aguirre von Wobeser E, Dekker HL, Huisman J, Matthijs HC. Time-series resolution of gradual nitrogen starvation and its impact on photosynthesis in the cyanobacterium Synechocystis PCC 6803. Physiol Plant. 2012;145:426–39.

[39] Llácer JL, Espinosa J, Castells MA, Contreras A, Forchhammer K, Rubio V. Structural basis for the regulation of NtcA-dependent transcription by proteins PipX and PII. Proceedings of the National Academy of Sciences of the United States of America. 2010;107:15397–402.

[40] Imamura S, Tanaka K, Shirai M, Asayama M. Growth phase-dependent activation of nitrogen-related genes by a control network of group 1 and group 2 sigma factors in a cyanobacterium. J Biol Chem. 2006;281:2668–75.

[41] Muro-Pastor AM, Herrero A, Flores E. Nitrogen-regulated group 2 sigma factor from Synechocystis sp. strain PCC 6803 involved in survival under nitrogen stress. J Bacteriol 2001;183:1090–5.

[42] Antal T, Kurkela J, Parikainen M, Kårlund A, Hakkila K, Tyystjärvi E, et al. Roles of group 2 sigma factors in acclimation of the cyanobacterium Synechocystis sp. PCC 6803 to nitrogen deficiency. Plant Cell Physio. 2016;57:1309–18.

[43] Osanai T, Imamura S, Asayama M, Shirai M, Suzuki I, Murata N, et al. Nitrogen Induction of Sugar Catabolic Gene Expression in Synechocystis sp. PCC 6803. DNA Research. 2006;13:185–95.

[44] Aguirre von Wobeser E, Ibelings BW, Bok J, Krasikov V, Huisman J, Matthijs HCP. Concerted Changes in Gene Expression and Cell Physiology of the Cyanobacterium Synechocystis sp. Strain PCC 6803 during Transitions between Nitrogen and Light-Limited Growth. Plant Physiology. 2011;155:1445–57.

[45] Klotz A, Georg J, Bucinska L, Watanabe S, Reimann V, Januszewski W, et al. Awakening of a Dormant Cyanobacterium from Nitrogen Chlorosis Reveals a Genetically Determined Program. Curr Biol. 2016;26:2862–72.

[46] Steuten B, Hoch PG, Damm K, Schneider S, Köhler K, Wagner R, et al. Regulation of transcription by 6S RNAs: Insights from the Escherichia coli and Bacillus subtilis model systems. RNA Biology. 2014;11:508–21.

[47] Sauer J, Schreiber U, Schmid R, Volker U, Forchhammer K. Nitrogen starvation-induced chlorosis in Synechococcus PCC 7942. Low-level photosynthesis as a mechanism of long-term survival. Plant Physiol. 2001;126:233–43.

[48] de Porcellinis AJ, Klahn S, Rosgaard L, Kirsch R, Gutekunst K, Georg J, et al. The Non-Coding RNA Ncr0700/PmgR1 is Required for Photomixotrophic Growth and the Regulation of Glycogen Accumulation in the Cyanobacterium Synechocystis sp. PCC 6803. Plant Cell Physiol. 2016;57:2091–103.

[49] Klähn S, Schaal C, Georg J, Baumgartner D, Knippen G, Hagemann M, et al. The sRNA NsiR4 is involved in nitrogen assimilation control in cyanobacteria by targeting glutamine synthetase inactivating factor IF7. Proc Natl Acad Sci U S A. 2015;112:E6243–52.

[50] Mitschke J, Georg J, Scholz I, Sharma CM, Dienst D, Bantscheff J, et al. An experimentally anchored map of transcriptional start sites in the model cyanobacterium Synechocystis sp. PCC6803. Proc Natl Acad Sci U S A. 2011;108:2124–9.

[51] Koskinen S, Hakkila K, Gunnelius L, Kurkela J, Wada H, Tyystjärvi T. In vivo recruitment analysis and a mutant strain without any group 2 s factor reveal roles of different s factors in cyanobacteria. Molecular Microbiology. 2016;99:43–54.

[52] Schwarz R, Forchhammer K. Acclimation of unicellular cyanobacteria to macronutrient deficiency: emergence of a complex network of cellular responses. Microbiology. 2005;151:2503–14.

[53] Osanai T, Ikeuchi M, Tanaka K. Group 2 sigma factors in cyanobacteria. Physiol Plant. 2008;133:490–506.

[54] Los DA, Zorina A, Sinetova M, Kryazhov S, Mironov K, Zinchenko VV. Stress sensors and signal transducers in cyanobacteria. Sensors (Basel). 2010;10:2386–415.

[55] Kopf M, Klahn S, Scholz I, Matthiessen JK, Hess WR, Voss B. Comparative analysis of the primary transcriptome of Synechocystis sp. PCC 6803. DNA Res. 2014;21:527–39.

[56] Neusser T, Polen T, Geissen R, Wagner R. Depletion of the non-coding regulatory 6S RNA in E. coli causes a surprising reduction in the expression of the translation machinery. BMC Genomics. 2010;11:165-.

[57] Trotochaud AE, Wassarman KM. A highly conserved 6S RNA structure is required for regulation of transcription. Nat Struct Mol Biol 2005;12:313–9.

[58] Beckmann BM, Burenina OY, Hoch PG, Kubareva EA, Sharma CM, Hartmann RK. In vivo and in vitro analysis of 6S RNA-templated short transcripts in Bacillus subtilis. RNA Biology. 2011;8:839–49.

[59] Asayama MIS, Yoshihara S, Miyazaki A, Yoshida N, Sazuka T, Kaneko T, Ohara O, Tabata S, Osanai T., SigC, the group 2 sigma factor of RNA polymerase, contributes to the late-stage gene expression and nitrogen promoter recognition in the cyanobacterium Synechocystis sp. strain PCC 6803. Biosci Biotechnol Biochem. 2004;68:477–87.

[60] Tuominen I, Pollari M, Tyystjärvi E, Tyystjärvi T. The SigB σ factor mediates high-temperature responses in the cyanobacterium Synechocystis sp. PCC6803. FEBS Letters. 2006;580:319–23.

[61] Pollari M, Gunnelius L, Tuominen I, Ruotsalainen V, Tyystjärvi E, Salminen T, et al. Characterization of Single and Double Inactivation Strains Reveals New Physiological Roles for Group 2 s Factors in the Cyanobacterium Synechocystis sp. PCC 6803. Plant Physiology. 2008;147:1994–2005.

[62] Nikkinen H-L, Hakkila K, Gunnelius L, Huokko T, Pollari M, Tyystjärvi T. The SigB s Factor Regulates Multiple Salt Acclimation Responses of the Cyanobacterium Synechocystis sp. PCC 6803. Plant Physiology. 2012;158:514–23.

[63] Tuominen I, Tyystjärvi E, Tyystjärvi T. Expression of Primary Sigma Factor (PSF) and PSF-Like Sigma Factors in the Cyanobacterium Synechocystis sp. Strain PCC 6803. Journal of Bacteriology. 2003;185:1116–9.

[64] Nakaya Y, Iijima H, Takanobu J, Watanabe A, Hirai MY, Osanai T. One day of nitrogen starvation reveals the effect of sigE and rre37 overexpression on the expression of genes related to carbon and nitrogen metabolism in Synechocystis sp. PCC 6803. Journal of Bioscience and Bioengineering. 2015;120:128–34.

[65] Osanai T, Kanesaki Y, Nakano T, Takahashi H, Asayama M, Shirai M, et al. Positive Regulation of Sugar Catabolic Pathways in the Cyanobacterium Synechocystis sp. PCC 6803 by the Group 2 σ Factor SigE. Journal of Biological Chemistry. 2005;280:30653–9.

[66] Osanai T, Imashimizu M, Seki A, Sato S, Tabata S, Imamura S, et al. ChlH, the H subunit of the Mg-chelatase, is an anti-sigma factor for SigE in Synechocystis sp. PCC 6803. Proceedings of the National Academy of Sciences of the United States of America. 2009;106:6860–5.

[67] Trautmann D, Voss B, Wilde A, Al-Babili S, Hess WR. Microevolution in cyanobacteria: Re-sequencing a motile substrain of Synechocystis sp. PCC 6803. DNA Res. 2012;19:435–48.

[68] Rippka R, Deruelles J, Waterbury JB, Herdman M, Stanier RY. Generic assignments, strain histories and properties of pure cultures of cyanobacteria. J Gen Microbiol. 1979;111:1–61.

[69] Zinchenko VV, Piven, I. V., Melnik, V. A. & Shestakov, S. V. Vectors for the complementation analysis of cyanobacterial mutants. Russ J Genet. 1999;35:228–32.

[70] Tandeau de Marsac N, Houmard J. Complementary chromatic adaptation: Physiological conditions and action spectra. Methods Enzymol 1988;167:318–28.

[71] Gründel M, Scheunemann R, Lockau W, Zilliges Y. Impaired glycogen synthesis causes metabolic overflow reactions and affects stress responses in the cyanobacterium Synechocystis sp. PCC 6803. Microbiology. 2012;158:3032–43.

[72] Gunnelius L, Tuominen I, Rantamäki S, Pollari M, Ruotsalainen V, Tyystjärvi E, et al. SigC sigma factor is involved in acclimation to low inorganic carbon at high temperature in Synechocystis sp. PCC 6803. Microbiology. 2010;156:220–9.

[73] Gunnelius L, Hakkila K, Kurkela J, Wada H, Tyystjärvi E, Tyystjärvi T. The omega subunit of the RNA polymerase core directs transcription efficiency in cyanobacteria. Nucleic Acids Research. 2014;42:4606–14.

[74] Dienst D, Duhring U, Mollenkopf HJ, Vogel J, Golecki J, Hess WR, et al. The cyanobacterial homologue of the RNA chaperone Hfq is essential for motility of Synechocystis sp. PCC 6803. Microbiology. 2008;154:3134–43.

[75] Dienst D, Georg J, Abts T, Jakorew L, Kuchmina E, Borner T, et al. Transcriptomic response to prolonged ethanol production in the cyanobacterium Synechocystis sp. PCC6803. Biotechnol Biofuels. 2014;7:21.

[76] Ritchie ME, Phipson B, Wu D, Hu Y, Law CW, Shi W, et al. limma powers differential expression analyses for RNA-sequencing and microarray studies. Nucleic Acids Research. 2015;43:e47–e.

